# Prefrontal oscillations modulate the propagation of neuronal activity required for working memory

**DOI:** 10.1101/531574

**Authors:** Jason S. Sherfey, Salva Ardid, Earl K. Miller, Michael E. Hasselmo, Nancy J. Kopell

## Abstract

Cognition involves using attended information, maintained in working memory (WM), to guide action. During a cognitive task, a correct response requires flexible, selective gating so that only the appropriate information flows from WM to downstream effectors that carry out the response. In this work, we used biophysically-detailed modeling to explore the hypothesis that network oscillations in prefrontal cortex (PFC), leveraging local inhibition, can independently gate responses to items in WM. The key role of local inhibition was to control the period between spike bursts in the outputs, and to produce an oscillatory response no matter whether the WM item was maintained in an asynchronous or oscillatory state. We found that the WM item that induced an oscillatory population response in the PFC output layer with the shortest period between spike bursts was most reliably propagated. The network resonant frequency (i.e., the input frequency that produces the largest response) of the output layer can be flexibly tuned by varying the excitability of deep layer principal cells. Our model suggests that experimentally-observed modulation of PFC beta-frequency (15-30 Hz) and gamma-frequency (30-80 Hz) oscillations could leverage network resonance and local inhibition to govern the flexible routing of signals in service to cognitive processes like gating outputs from working memory and the selection of rule-based actions. Importantly, we show for the first time that nonspecific changes in deep layer excitability can tune the output gate’s resonant frequency, enabling the specific selection of signals encoded by populations in asynchronous or fast oscillatory states. More generally, this represents a dynamic mechanism by which adjusting network excitability can govern the propagation of asynchronous and oscillatory signals throughout neocortex.

## Introduction

Many cognitive tasks require flexible routing of information from multiple sources to update internal rule representations and guide responses (***Melrose et al., 2007***; ***Badre and Frank, 2012***; ***Buschman et al., 2012***; ***Hasselmo and Stern, 2018***; ***Bhandari and Badre, 2018***; ***Zhu et al., 2018***, ***2020***). Cognition involves routing of attended information to guide correct responses based on information that has been buffered in working memory (WM) (***Baddeley and Hitch, 1974***; ***Miller, 2000***). The routing must regulate the flow of information about sensory stimuli, about rules to be applied and feedback about responses.

Many experiments demonstrate that all facets of cognition are associated with the modulation of oscillatory dynamics in networks of neurons (***Cho et al., 2006***; ***Tzur and Berger, 2009***; ***Siegel et al., 2009***; ***Buschman et al., 2012***; ***Brincat and Miller, 2016***; ***Lundqvist et al., 2016***, ***2018***). Much discussion has addressed the role of oscillatory dynamics versus neuron spike rates in routing cognitively-relevant representations (***Ardid et al., 2010***; ***Mante et al., 2013***; ***Ardid et al., 2019***). In this work, we investigate routing from the perspective of a network in which internal dynamics regulate the feedforward transmission (i.e., a relay network) that selectively propagates one or more competing signals, effectively gating out signals that lose the competition. Specifically, we used biophysically-detailed modeling to explore the hypothesis that network oscillations in the relay network, leveraging local inhibition, can independently gate responses to items represented in source populations like those encoding items in working memory.

Experiments and models have demonstrated that WM items in PFC may be represented in populations with spiking that is asynchronous (***Wang, 1999***; ***Tegnér et al., 2002***; ***Renart et al., 2010***) or synchronous with rhythmic modulation (***Compte et al., 2000***; ***Lundqvist et al., 2010***, ***2018***) over some interval of time. Such asynchronous and oscillatory population activities are indicative of local networks in different dynamical states (***Tegnér et al., 2002***; ***Akam and Kullmann, 2010***). Notably, modulation of beta (15-30 Hz) and gamma (30-80 Hz) frequency oscillations has been observed in PFC during WM tasks when items were loaded, maintained, and retrieved from WM (***Siegel et al., 2009***; ***Lundqvist et al., 2016***, ***2018***; ***Bastos et al., 2018***). ***Akam and Kullmann*** (***2010***) showed how a band-pass filter network based on resonant, feedforward inhibition supports the selective read-out of activity with gamma-frequency modulation. Their model was tuned to a balanced regime in which excitatory cells were unresponsive to asynchronous inputs. In contrast to the band-pass filter model, we focus on a network model with strong, feedback inhibition that exhibits resonance (i.e., larger responses to preferred input frequencies) while remaining responsive to asynchronous signals. This enabled us to consider under which conditions signals in an asynchronous or oscillatory state may be propagated downstream for retrieval from WM.

We have previously shown through detailed modeling that a PFC network with strong feedback inhibition exhibits resonance (***Sherfey et al., 2018a***). In that previous work, we presented a scenario where a resonant rhythm suppressed an asynchronous signal. In the results presented here we investigated competition between asynchronous, high beta-frequency (20-30 Hz), and gamma-frequency rhythmic input signals. Most importantly, we show how the output network can be flexibly tuned to selectively propagate signals in any of these states while blocking the others through biased competition. Throughout this paper, we will show how this behavior enables the deep layers of PFC to function as an output gate and consider implications for WM. We found that, given multiple excitatory populations in an output layer, whichever output population has the shortest period between spike volleys (i.e., bursts of action potentials) most reliably engages local inhibition which, in turn, suppresses responses in all opposing populations. We show that asynchronous inputs and beta-rhythmic inputs can lead to a higher output frequency than a faster gamma-rhythmic input. Importantly, we show for the first time that nonspecific changes in deep layer excitability can tune the output gate’s resonant frequency, enabling the specific selection of signals encoded by populations in asynchronous or fast oscillatory states. In our PFC model, these dynamics yield a flexible mechanism for gating the flow of information.

This paper begins with the details of our computational model followed by a simulation study to investigate the mechanisms that underlie oscillatory output gating, their generality, and the ability to flexibly tune the gate to select input items from populations in different dynamical states. Gating rules and control mechanisms will be summarized in the Discussion, and the paper will close with a detailed look at how oscillatory gating could serve working memory and cognitive control.

## Methods

### Cortical network models of the output gate

The network model represents a cortical output layer with excitatory principal cells (PCs) connected reciprocally to inhibitory interneurons (INs). Hodgkin-Huxley type PC and IN models were taken from a computational representation of a deep layer PFC network consisting of two-compartment PCs (soma and dendrite) with ion channels producing *I*_*NaF*_, *I*_*KDR*_, *I*_*NaP*_, *I*_*Ks*_, *I*_*Ca*_, and *I*_*KCa*_ currents (μA/cm^2^) and fast spiking INs with channels producing *I*_*NaF*_ and *I*_*KDR*_ currents (***Durstewitz and Seamans, 2002***) (***Figure 1***A; see figure caption for channel definitions). IN cells had spike-generating *I*_*NaF*_ and *I*_*KDR*_ currents with more hyperpolarized kinetics and faster sodium inactivation than PCs, resulting in a more excitable interneuron with fast spiking behavior. In the baseline case, PC and IN cell models were identical to those in the original published work while network connectivity was adjusted to produce natural oscillations (not in ***Durstewitz and Seamans*** (***2002***)), as described below. All cells were modeled using a conductance-based framework with passive and active electrical properties of the soma and dendrite constrained by experimental considerations (***Durstewitz et al., 2000***). Membrane potential *V* (mV) was governed by:

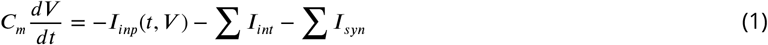

where *t* is time (ms), *C*_*m*_ = 1 μF/cm^2^ is the membrane capacitance, *I*_*int*_ denotes the intrinsic membrane currents (μA/cm^2^) listed above, *I*_*inp*_(*t, V*) is an excitatory current (μA/cm^2^) reflecting inputs from external sources described below, and *I*_*syn*_ denotes synaptic currents (μA/cm^2^) driven by PC and IN cells in the network. This model represents a WM-related prefrontal network because it was constrained by in vitro data from PFC (***Durstewitz et al., 2000***) in neurons that exhibit physiology (e.g., delay activity) known to support WM functions (***Seamans et al., 2008***). Furthermore, the inputs and network connectivity (described below) reflect the feedforward, interlaminar projections and interneuron-mediated inhibition observed in PFC (***Kritzer and Goldman-Rakic, 1995***; ***DeFelipe, 1997***) and other cortical regions (***Douglas and Martin, 2004***).

**Figure 1.**
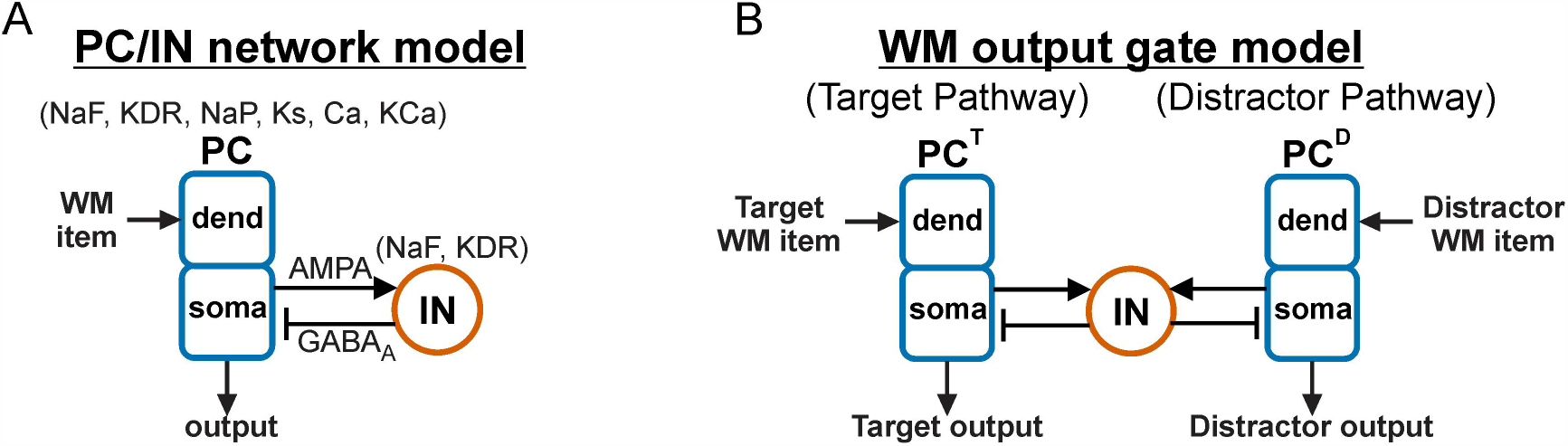
Architecture of output networks. (**A**) Diagram showing feedforward excitation from external independent Poisson spike trains to the dendrites of 20 two-compartment (soma, dend) principal cells (PCs) receiving feedback inhibition from a population of 5 fast spiking interneurons (INs). All PC and IN cells have biophysics based on prefrontal neurons (***Durstewitz and Seamans, 2002***) (Ion channel key: NaF = fast sodium channel; KDR = fast delayed rectifier potassium channel; NaP = persistent sodium channel; Ks = slow (M-type) potassium channel; Ca = high-threshold calcium channel; KCa = calcium-dependent potassium channel). (**B**) Diagram showing a rhythmically-driven target population of PCs (PC^T^) competing with an asynchronously-driven distractor population (PC^D^) through a shared population of inhibitory IN cells. This figure was adapted from ***Sherfey et al.*** (***2018a***).

The output layer had either one or two populations of PCs with each output population receiving inputs from one or two input signals that we will interpret as encoding working memory (WM) items (defined below) to illustrate the utility of oscillatory gating. Natural and resonant properties were characterized using a network with one input item and one output population (***Figure 1***A). Competition was investigated using a network with one or two homogeneous PC populations driven by multiple input items while interacting through a shared population of inhibitory cells (***Figure 1***B).

### Model complexity

We investigated both larger-scale, more-detailed and simplified versions of the model. The more-detailed model had a comparable number of neurons as in ***Durstewitz et al.*** (***2000***) (100 PCs and 25 INs) or 10 times as many.

The simplified model had 20 PCs, 5 INs and less detailed connectivity (described below). We first determined that biased competition behaved similarly in the two models (see Results). Then, we used the simplified model, exhibiting the same essential behavior, to explore the effects of competition over a larger region of parameter space. This exploration involved > 50,000 simulations to investigate dependence of network resonance on input properties in ***Sherfey et al.*** (***2018a***); > 50,000 to identify regions of interest in parameter space for this work on competition-based gating; and > 5,000 simulations for figures. Most figures show results for the simplified model to maintain consistency of presentation throughout the manuscript; however, it will be shown that the detailed and simplified models exhibit qualitatively similar gating phenomena (see ***Figure 2***). Furthermore, a much simpler network model of leaky integrate-and-fire (LIF) neurons reproduces the qualitative behavior (Supplementary Figure 1) and could be used for further study of generic network dynamics. A generic LIF network would lack specificity of prefrontal conductances of the model explored in the present work.

**Figure 2.**
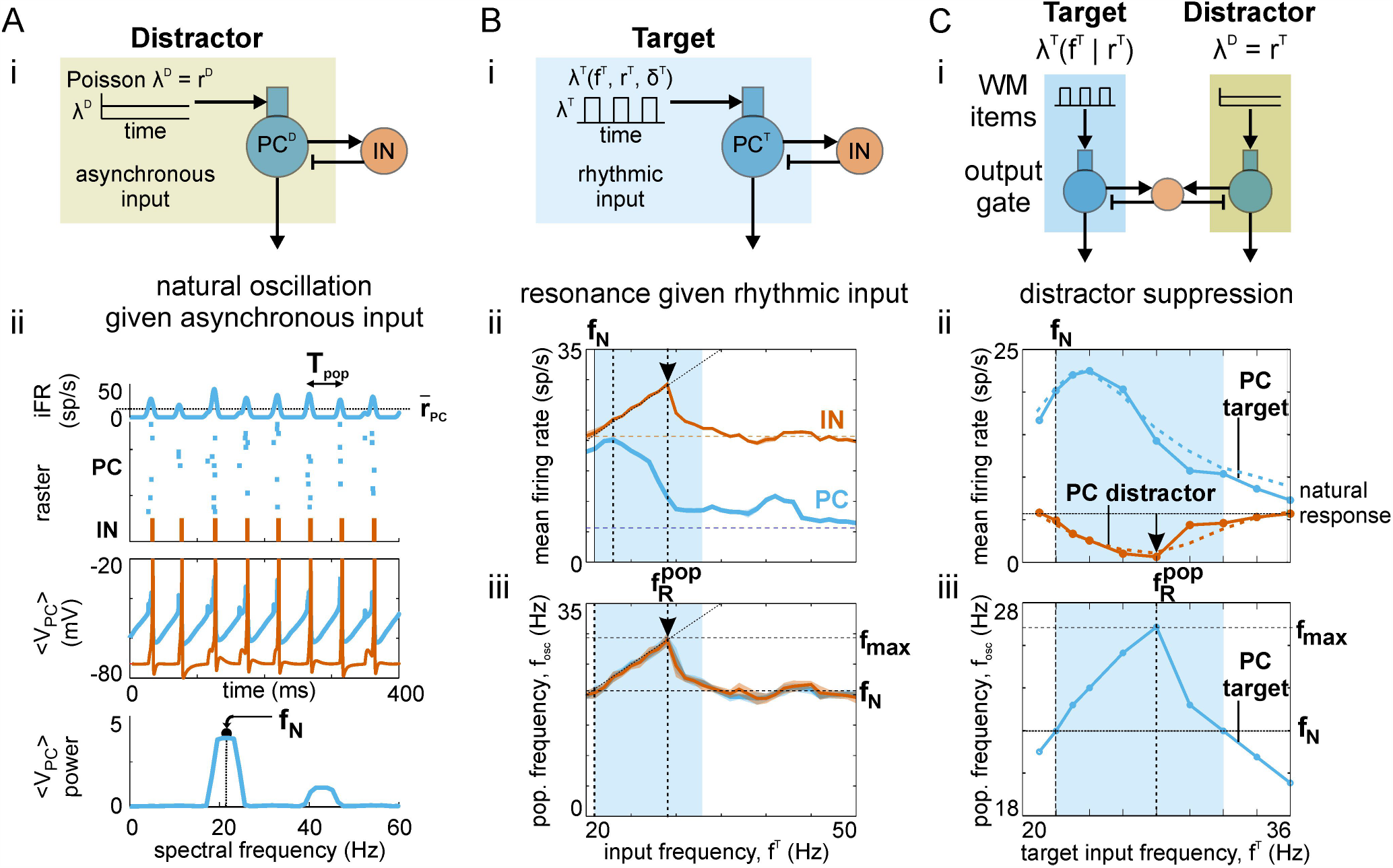
Dynamics of PC/IN network. (**A**) Natural oscillation. (i) Schematic showing asynchronous Poisson spike trains driving the dendrites of two-compartment principal cells (PCs) coupled to interneurons (INs) providing strong feedback inhibition. (ii) The response of the asynchronously-driven PC/IN network is an inhibition-paced, pulsatile oscillation of periodic spike volleys occurring at a natural frequency, *f*_*N*_. (**B**) Resonant response. (i) Schematic showing rhythmically-modulated Poisson spike trains driving the PC/IN network. The input has square wave rate-modulation with a 10 ms pulse width; i.e., all input spikes occur within 10 ms on a given cycle. (ii) The impact of the frequency of input rate-modulation, *f*^*T*^, on mean output firing rates (averaged over time, population, and 10 realizations) for PC (blue) and IN (red) populations. (iii) The impact of the input frequency on the frequency of output rate-modulation, *f*_*pop*_, for PC (blue) and IN (red) populations. Vertical dashed lines mark the resonant input frequencies. Horizontal dashed lines indicate the natural response to asynchronous input. The interval with blue shading highlights the range of input frequencies that yield an output oscillation that is faster than the natural frequency. (**C**) Distractor suppression. (i) Schematic showing a rhythmically-driven target population of PCs (PC^T^) competing with an asynchronously-driven distractor population (PC^D^) through a shared pool of inhibitory INs. Strong feedback inhibition causes each output to be rhythmic; the shared INs induce competitive interactions between them. (ii) Mean firing rate outputs for target (blue) and distractor (red) as target input frequency is increased; results for detailed and simplified models are shown using dashed and solid lines, respectively. Peak target output, maximal distractor suppression, and the range of target input frequencies that suppressed the distractor response were the same in both models. Blue shading highlights the range of input frequencies yielding an output oscillation that is faster than the natural frequency; the range increases with input synchrony and strength (not shown). (iii) Output population frequency of the target (blue) and the natural frequency of an asynchronously-driven PC/IN network (horizontal dashed line). The vertical dashed line marks the peak output frequency of the target and corresponds to the target input frequency at which the distractor output is maximally suppressed. Parts of (A) and the simplified model in (C) were adapted from ***Sherfey et al.*** (***2018a***).

### Network connectivity

PCs provided excitation to IN cells mediated by *α*-amino-3-hydroxy-5-methyl-4-isoxazolepropionic acid (AMPA) currents. IN cells in turn provided strong feedback inhibition mediated by *γ*-aminobutyric acid (GABA_A_) currents to PCs. This combination of fast excitation and strong feedback inhibition is known to generate robust network oscillations in response to tonic drive (***Whittington et al., 2000***; ***Börgers and Kopell, 2005***). Connection probabilities in the detailed model were 40% from PC-to-IN and 60% from IN-to-PC. Connectivity was all-to-all in the simplified model to ensure that a small number of active neurons were sufficient to produce an ongoing network oscillation. AMPA currents were modeled by:

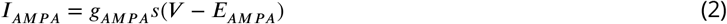

where *V* is the postsynaptic membrane voltage, *g*_*AMPA*_ is the maximal synaptic conductance, *s* is a synaptic gating variable, and *E*_*AMPA*_ = 0 mV is the synaptic reversal potential. Synaptic gating was modeled using a first-order kinetics scheme:

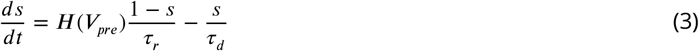

where *V*_*pre*_ is the presynaptic membrane voltage, *τ*_*r*_ = 0.4 ms and *τ*_*d*_ = 2 ms are time constants for neurotransmitter release and decay, respectively, and *H*(*V*) = (1 + *tanh*(*V*/4))/2 is a sigmoidal approximation to the Heaviside step function. In simulations with connectivity between PCs, AMPA synapses were modeled using the same kinetics. GABA_A_ currents were modeled in the same way with *E*_*GABA*_ = −75 mV and *τ*_*d*_ = 5 ms. Inhibition between IN cells was included in the detailed model; however, it was excluded from the simplified model since it did not affect the network dynamics as long as IN cells were synchronously activated by the PC population and inhibited each other with the same decay time constant. NMDA and GABA_B_ currents were excluded from both models. Maximum synaptic conductances for PCs were (in mS/cm^2^): AMPA (.03), GABA_A_ (.1); for IN cells: AMPA (.03), GABA_A_ (.1).

### Working memory items

Each PC in the output gate received independent Poisson spike trains with time-varying instantaneous rate *λ*(*t*) (sp/s) and time-averaged rate *r* = ⟨*λ*⟩. Spikes were integrated in a synapse *s*_*inp*_ with exponential AMPAergic decay contributing to an excitatory synaptic current *I*_*inp*_ = *g_inp_s_inp_*(*V* − *E_AMPA_*) with maximal conductance *g*_*inp*_ (mS/cm^2^). WM items were modeled by collections of spike trains with the same instantaneous rate-modulation. A given input item to a PC output population can be interpreted as conveying information from a population of excitatory pyramidal cells in superficial PFC that participate in a network in a particular dynamical state (i.e., asynchronous or oscillatory).

Population rate-coding was incorporated into a WM item using a spatial pattern of time-averaged firing rates *r*(*x*) for spike trains driving an output population with cells indexed by *x*. That is, output cell *x* was driven by a collection of spike trains with average firing rate *r*(*x*). Spatial profiles were either uniform (i.e., the average drive to each output cell had the same strength) or Gaussian bumps centered on particular output neurons. WM items encoded by populations in different dynamical states were generated by modulating instantaneous rates *λ*(*t*). Signals representing WM items in an asynchronous state were modeled by homogeneous Poisson spike trains with constant rate *λ*(*t*) = *r* whereas signals representing WM items in an oscillatory state were modeled using rate-modulated Poisson processes with periodically-modulated instantaneous rates. WM items with sine wave modulation had *λ*(*t*) = *r*(1 + *sin*(2*πft*))/2 parameterized by *r* (sp/s) and rate modulation frequency *f* (Hz). We also investigated oscillatory inputs with square wave modulation in order to differentiate the effects of synchrony and frequency while maintaining the ability to compare our results with other studies. In this work, we define synchrony operationally as the interval over which spikes are highly concentrated on an average cycle. Using a square wave oscillation enables the interval over which spikes are generated on a given cycle to be held constant while the frequency is varied. Square wave rate-modulation results in periodic trains of spikes (pulse packets) parameterized by *r* (sp/s), inter-pulse frequency *f* (Hz), and pulse width *δ* (ms). *δ* reflects the synchrony of spikes in the source population with smaller values implying greater synchrony; decreasing *δ* corresponds to decreasing the duty cycle of the square wave. For the square wave input, we chose to hold constant *r* so that across frequencies the only significant change is in the patterning of spikes and not the total number of spikes; this results in larger pulses being delivered to postsynaptic PCs at lower frequencies, consistent with lower frequencies engaging larger networks (***Nunez et al., 2006***). More advanced approaches to generating periodic, quasi-periodic, and non-periodic synchronous spike trains exist (***Brette and Guigon, 2003***) but are not necessary to examine the effects of synchrony on biased competition in the networks studied here. Throughout the paper, WM item parameters *r*, *f*, and *δ* are assigned superscripts *T* or *D* indicating whether they represent “target” or “distractor” items, respectively (see ***Table 1*** for notation details).

**Table 1.** Meaning of symbols used in the study of output gating for working memory.

| Symbol | Description |
| --- | --- |
| **Parameters used to define working memory (WM) items** |  |
| $\lambda^T = \lambda^T(t)$ | Instantaneous rate of Poisson process for a Target item (kHz) |
| $r^T$ | Target item strength, $\langle \lambda^T \rangle_t$ (kHz) |
| $f^T$ | Target item oscillation frequency of rate-modulation (Hz) |
| $\delta^T$ | Target item synchrony control parameter; pulse width of square wave rate-modulation (ms) |
| $\lambda^D = \lambda^D(t)$ | Instantaneous rate of Poisson process for a Distractor item (kHz) |
| $r^D$ | Distractor item strength, $\langle \lambda^D \rangle_t$ (kHz) |
| $f^D$ | Distractor item oscillation frequency of rate-modulation (oscillatory distractors only) (Hz) |
| **Output gate** |  |
| $PC^T$ | Principal cell population driven by a Target item |
| $PC^D$ | Principal cell population driven by a Distractor item |
| $IN$ | Inhibitory interneurons providing strong feedback and lateral inhibition |
| $\bar{r}_{PC}^T$ | Target output strength; time-averaged population firing rate of $PC^T$ (sp/s) |
| $\bar{r}_{PC}^D$ | Distractor output strength; time-averaged population firing rate of $PC^D$ (sp/s) |
| $\bar{r}_{IN}$ | Time-averaged population firing rate of interneurons (sp/s) |
| $f_{pop}^T$ | Target output oscillation frequency (inverse of period between $PC^T$ spike volleys) (Hz) |
| $f_{pop}^D$ | Distractor output oscillation frequency (inverse of period between $PC^D$ spike volleys) (Hz) |
| $f_N$ | Natural oscillation frequency of asynchronously-driven output gate (Hz) |
| $f_{max}$ | Maximum oscillation frequency of rhythmically-driven output gate (Hz) |
| $f_R^{pop}$ | $f_{pop}$-resonant frequency of output gate (Hz) (i.e., item $f^T$ that yields $f_{pop}^T = f_{max}$) |

All principal cells in the output gate received additional asynchronous inputs representing uncorrelated background activity from 100 cells in other brain areas spiking at 1 sp/s. Notably, feedforward inhibition was excluded from the present work so that all inputs (i.e., asynchronous and oscillatory) were maximally effective at driving PCs in the output layer. Control values for WM item parameters were *r* = 1000 sp/s (corresponding to an input population with 1000 neurons spiking at 1 sp/s); *δ* = 1 ms (high synchrony), 10 ms (medium synchrony), or 19 ms (low synchrony), and *g*_*inp*_ = 0.0015 mS/cm^2^.

In simulations probing resonant properties of the output gate, the item modulation frequency *f* was varied from 20 Hz to 50 Hz (in 1 Hz steps) across simulations. In simulations exploring output gating among parallel pathways, input items had the same mean strength (i.e., *r*); this ensures that any difference between the ability of WM items to drive their corresponding outputs resulted from differences in dynamical states and not differences in average activity levels.

### Data analysis

For each simulation, instantaneous output firing rates, iFR, were computed with Gaussian kernel regression on population spike trains using a kernel with 6 ms width for visualization and 2 ms for calculating the power spectrum. Mean population firing rates, 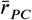 and 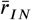, were computed by averaging iFR over time for PC and IN populations, respectively; they index overall activity levels by the average firing rate of the average cell in the population. The collective oscillation frequency of an output population, *f*_*pop*_, is defined as the dominant frequency of its iFR and was identified as the spectral frequency with peak power in the iFR spectrum (see ***Sherfey et al.*** (***2018a***) for more details). The natural frequency *f*_*N*_ of the output network was identified as the population frequency *f*_*pop*_ produced in response to an asynchronous input for a given input strength *r*.

Across simulations varying input frequencies, statistics were plotted as the mean ± standard deviation calculated across 10 realizations. The *f*_*pop*_-resonant frequency, 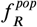, was defined as the input frequency that maximized output oscillation frequency *f*_*pop*_ in the beta/gamma range. We will show that maximizing *f*_*pop*_ is more important for biased competition in the relay network than maximizing output strength. Our measures of spiking and oscillatory activity in the strongly-driven network differs from measures used in work on resonance in weakly-driven networks (***Richardson et al., 2003***; ***Akam and Kullmann, 2010***; ***Ledoux and Brunel, 2011***) and is more similar to measures used for strongly-driven single cells (***Rotstein, 2017***). See ***Sherfey et al.*** (***2018a***) for a comparison of perspectives on resonance in cells and networks.

### Simulation tools

All models were implemented in Matlab using the DynaSim toolbox (***Sherfey et al., 2018b***) (http://dynasimtoolbox.org) and are publicly available online at: http://github.com/jsherfey/PFC_models. Numerical integration was performed using a 4th-order Runge-Kutta method with a fixed time step of 0.01 ms. Simulations were run for 2500 ms and repeated 10 times. The network was allowed to settle to steady-state before external signals were delivered at 400 ms. Plots of instantaneous responses begin at signal onset. The first 500 ms of response was excluded from analysis, although including the transient did not alter our results significantly.

## Results

This section begins with a review of relevant work on PFC network dynamics and an investigation of the mechanisms that underlie oscillatory gating mediated by biased competition. After that we will address how the gate can be tuned to select working memory (WM) items that are asynchronous or oscillatory with different frequencies. Finally, we will demonstrate the utility of our gating mechanism for winner-take-all selection of WM items delivered to the output gate along either parallel or convergent pathways.

### Background: inhibition-based oscillations and resonance in cortical networks

As was mentioned above, high beta (20-30Hz, which we will call “beta” for simplicity) and gamma (30-80Hz) oscillations are modulated in PFC during *in vivo* WM tasks (***Cho et al., 2006***; ***Tzur and Berger, 2009***; ***Siegel et al., 2009***; ***Lundqvist et al., 2016***, ***2018***; ***Bastos et al., 2018***). Neuronal networks generate oscillations at these frequencies given tonic drive (e.g., steady asynchronous spiking, current injection, or a glutamate receptor agonist) and strong feedback inhibition with inhibitory time constants comparable to those measured from the IPSCs produced by fast spiking, parvalbumin-positive interneurons in PFC (***Adams et al., 2017***; ***Sherfey et al., 2018a***). Such inhibition-based oscillations (***Whittington et al., 2000***) output periodic volleys of spikes (***Figure 2***A, raster plot) and generate a population frequency *f*_*pop*_ (i.e., the inverse of the period between spike volleys) that depends on the strength of input drive. See Methods for how we compute *f*_*pop*_ and ***Table 1*** for a summary of all symbols used throughout the paper. The population frequency in response to an asynchronous drive will be called the “natural frequency”, *f*_*N*_, of the network; in this case, *f*_*N*_ = 21 Hz. We will demonstrate the importance of the natural frequency for determining responses in an output gate below.

Since we are interested in gating the propagation of signals reflecting different dynamical states, perhaps encoding WM items, we will also review the impact of oscillatory inputs on PC/IN networks with one PC population (***Figure 2***B) before examining the competitive, output gate model. A rhythmically-driven PC population (***Figure 2***Bi) will produce spike volleys with a period matched to the input at low frequencies. At a sufficiently high input frequency, the depolarizing conductances in a fraction of PCs will fail to reach threshold on each cycle of the input; thus, the number of spikes per volley (i.e., the time-averaged firing rate) will decrease (***Figure 2***Bii). For a finite range of increasing frequencies, the output frequency of the PC/IN network will remain equal to the input frequency as long as a sufficient fraction of PCs are able to activate INs on every cycle of the input (***Figure 2***Biii). In this case, the output frequency peaks at *f*_*max*_ ≈ 28 Hz; however, ***Sherfey et al.*** (***2018a***) shows that *f*_*max*_ increases with the synchrony and strength of the oscillatory input. For much higher input frequencies ≫ *f*_*max*_, the oscillatory input acts like a tonic input, yielding an output with spike volleys occurring at the natural frequency of the network. Therefore, there is only a finite range of input frequencies for which the PC/IN network generates an output faster than the natural frequency (i.e., *f*_*pop*_ > *f*_*N*_) (***Figure 2***Biii, shaded region). This intermediate range of fast oscillation frequencies will be shown to impart to oscillatory inputs a competitive advantage in the output gate.

### Competition in the output gate

The more-detailed and simplified output gate models each consist of two PC populations connected to a common set of inhibitory INs that strongly inhibit the entire layer as described in Methods (***Figure 2***C). To begin, each PC population was driven by a distinct input, which we will call a WM item. Spiking in a given output population relays information from its source population in the unmodeled input layer, which we will call a WM buffer. We are interested in how the difference in spiking between the output populations depends on the oscillatory state of the WM items driving them; that is, how the dynamical state of items in the WM buffer affects the relative relay of information from the buffer. In some cases, all items will be relayed; in others, only a subset will be relayed. To describe our results, we will start by considering one input to be an oscillatory “target” item and another to be an asynchronous “distractor” item. We will examine under what conditions an oscillatory target will be propagated while suppressing the response to distractor through interneuron-mediated inhibition (***Figure 2***Ci). While there is accumulating evidence that WM item-encoding populations are oscillatory, different studies suggest different frequencies may be task-relevant (***Siegel et al., 2009***; ***Lundqvist et al., 2016***). Thus, we subsequently examine under what conditions different oscillatory signals (target and distractor) at different frequencies may be selectively propagated in the presence of multiple oscillations.

In the output gate model, the response to an asynchronous item (the distractor) was suppressed for a finite range of target frequencies, *f*^*T*^ (***Figure 2***Cii, shaded region). In fact, it was precisely the intermediate range of target input frequencies that yielded an output frequency, *f*_*pop*_, greater than the natural frequency (i.e., when target *f*_*pop*_ > *f*_*N*_) (***Figure 2***Ciii, compare solid and dashed lines; ***Figure 2***Cii-iii, compare shaded regions). Biased competition resulted in similar behavior in both the more-detailed and simplified models (compare solid and dashed lines in ***Figure 2***Cii). Maximum distractor suppression occurred at peak *f*_*pop*_ (***Figure 2***Cii-iii, vertical dashed lines) (i.e., when target *f*_*pop*_ = *f*_*max*_) and not at peak target spiking (***Figure 2***Cii-iii, compare blue target curves). This implies that a *f*_*pop*_-resonant item (i.e., an item that maximizes output population frequency) in a WM buffer will maximally suppress distractor responses in the output gate. Using the more-detailed model, ***Figure 3*** illustrates that the resonant bias enables items held in working memory by different populations, possibly in superficial layers of PFC, to be gated by the resonant properties of an output network, possibly in the deep layers of PFC. From a cognitive perspective, this suggests if superficial items represent different dimensions of a stimulus (S) and deep layer outputs map onto alternative action plans (R1, R2), then context-dependent rhythmicity can govern rule-based selection of stimulus-response mapping (see Discussion for further consideration). From the perspective of network competition, ***Figure 3*** shows that the resonance of the output network results in propagation of the oscillatory target input, and suppression of activity from the asynchronous distractor input.

**Figure 3.**
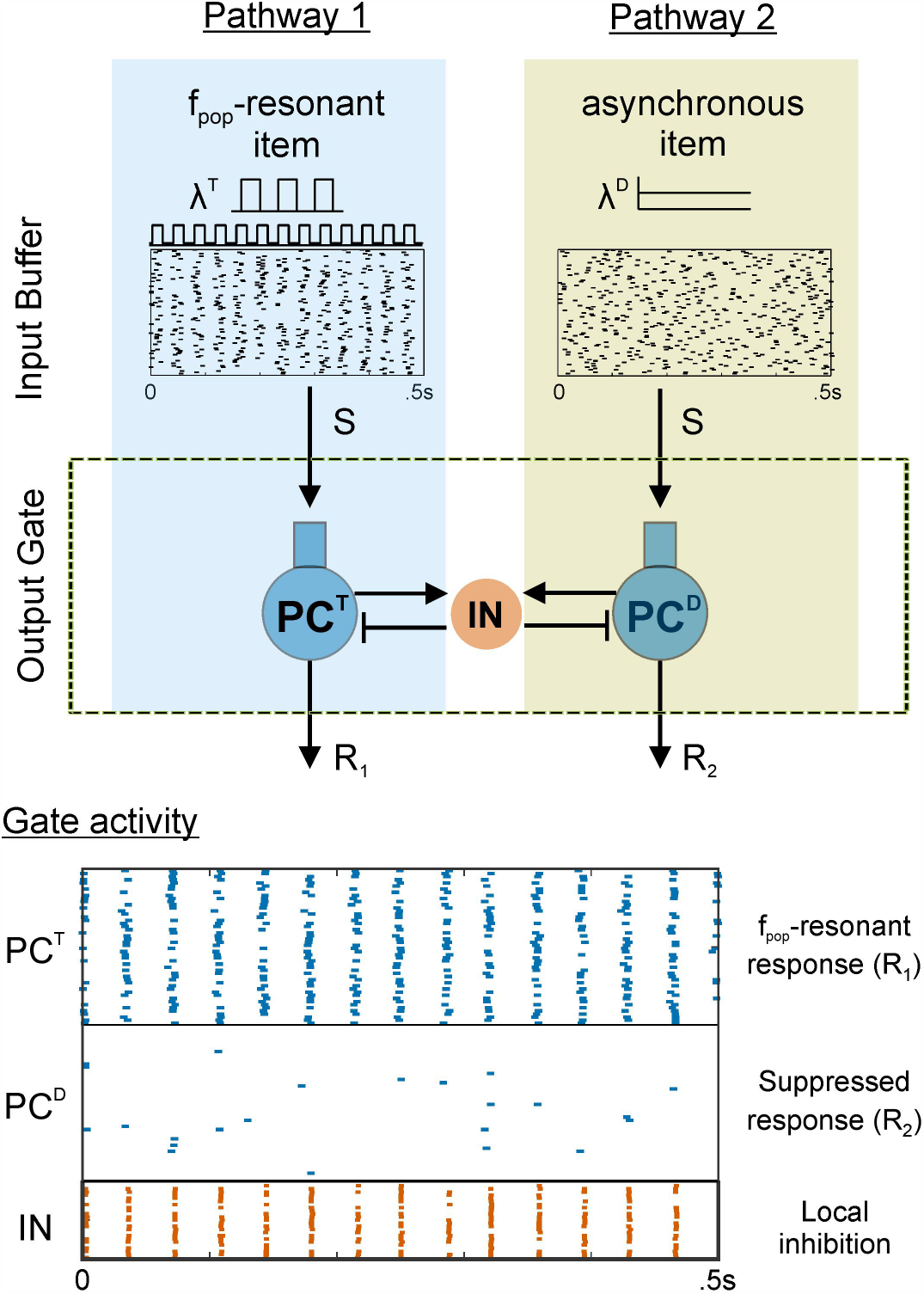
Resonant bias supports rule-based stimulus-response mapping. Example simulation of the more-detailed model showing two pathways (i.e., alternative stimulus-response mappings) with superficial layer inputs and deep layer outputs. One pathway has a resonant input that drives its output population while the other pathway has an asynchronous input and an output that is suppressed by interneuron-mediated lateral inhibition.

This oscillation-dependent, gating phenomenon can be understood most easily by considering the interaction between excitation and inhibition in the output gate over time (***Figure 4***). Essentially, the excitatory output gate population with the shortest period between spike volleys will be the dominant driver of local inhibition that suppresses spikes in all other populations connected to the same interneurons. The number of spikes per PC volley (i.e., the time-averaged PC firing rate) is less important than the period between volleys (i.e., the population frequency) as long as there are enough spikes in an excitatory volley to engage the inhibition.

**Figure 4.**
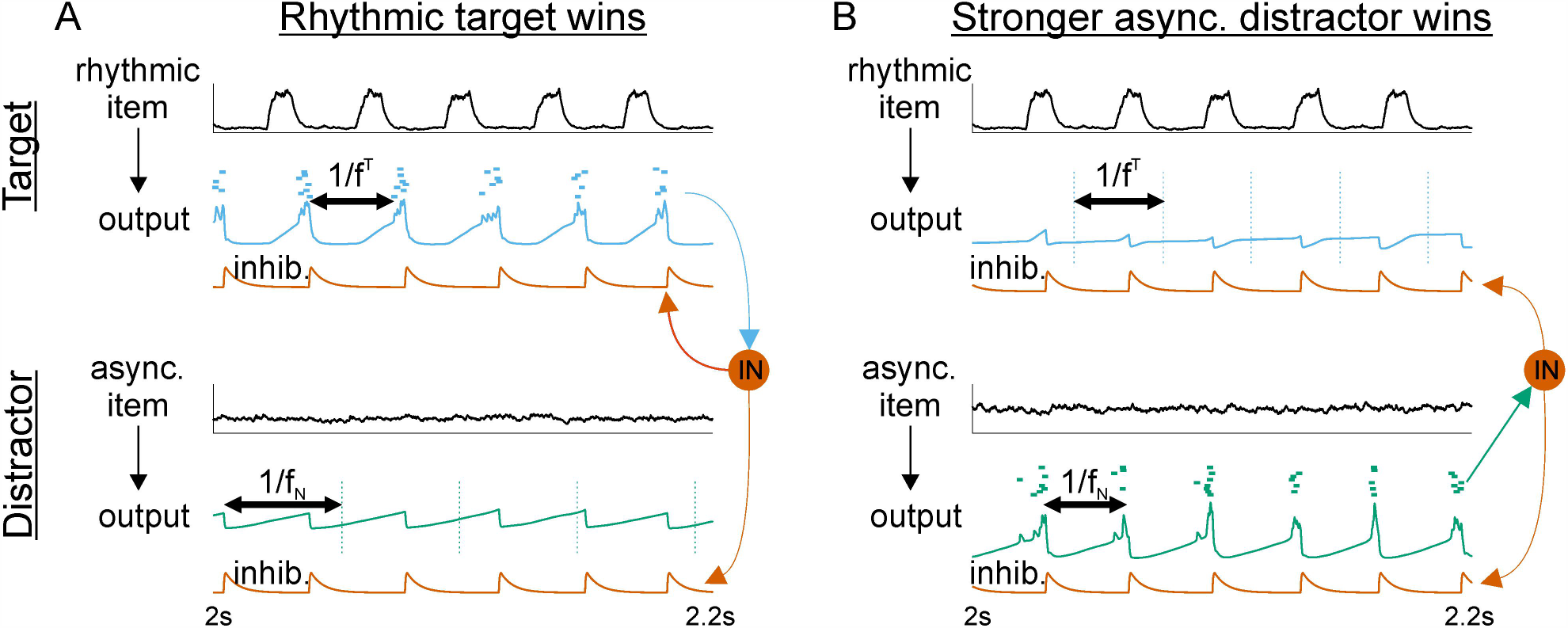
Oscillatory gating: control of periodic inhibition determines gate output. (**A**) Rhythmic WM item dominates the output gate. The response to the oscillatory input has a shorter period than the natural response to the asynchronous item (compare horizontal double arrows). Consequently, the target engages local inhibition to suppress the distractor response before it can reach threshold. Black curves show the excitatory postsynaptic potentials produced by (top) a rhythmic input item and (bottom) an asynchronous input item. Summed voltages and spikes for the (top) target and (bottom) distractor PC populations are shown beneath their inputs. Red curves show the inhibitory postsynaptic potentials produced by the INs onto (top) target and (bottom) distractor PCs. Vertical dashed lines on distractor output mark when spikes would have occurred in absence of lateral inhibition. (**B**) A stronger asynchronous WM item dominates the output gate. The response to a stronger asynchronous input produces a natural oscillation with a shorter period than that of the oscillatory input. Consequently, the distractor output engages local inhibition and suppresses the response to the oscillatory item. Vertical dashed lines on target output mark when spikes would have occurred in absence of lateral inhibition.

***Figure 4***A shows that when a periodic item in the WM buffer oscillated with frequency *f*_*N*_ < *f*^*T*^ < *f*_*max*_, its target output engaged interneurons in the output gate every 1/*f*^*T*^ seconds. Thus, the target output was dominant because it had a period shorter than the 1/*f*_*N*_ seconds required for the asynchronously-driven distractor output to reach threshold (***Figure 4***A; compare the periods marked by horizontal double arrows). We know from previous work (***Sherfey et al., 2018a***) that the natural frequency increases with the strength of an asynchronous drive. ***Figure 4***B shows that once an asynchronous item is strong enough for *f*_*N*_ to exceed *f*^*T*^, then the distractor output becomes the dominant driver of local inhibition in the output gate. Consequently, the response to an oscillatory item can be suppressed by a stronger asynchronous item when the latter induces a faster population frequency.

### Constraints on the suppression of stronger distractors

The previous simulations have demonstrated that either the oscillatory or asynchronous WM item can produce a larger response in the output gate PCs driven by them depending on the relative oscillation frequencies that they induce. Experiments have shown that task-relevant items in WM show context-dependent synchrony (***Buschman et al., 2012***) and that the strength of items may relate to stimulus familiarity (***Shen et al., 2018***). With this motivation, we sought next to investigate how the output gate response depends on the synchrony of oscillatory input items and to quantify how much of an advantage is provided by the oscillatory state.

We addressed this by delivering spikes from a 28Hz resonant, target item to one PC population in the output gate and spikes from an asynchronous, distractor item to a competing PC population in the output gate (***Figure 5***A). The strength of the distractor item, *r^D^*, was increased across simulations from 1.0 up to 2.0 times that of the target item. We found that the target output produced more spikes than the distractor output until the distractor item was 50% stronger than the target item (***Figure 5***B, green curve). Similar to above, the reason why the dominant output (i.e., the output producing the most spikes) switched at that point can be understood by considering the relative population frequencies of the two output populations. In the simulations here, the natural frequency of the distractor equals the resonant frequency of the target when the distractor item is 50% stronger. Across the set of simulations, whichever output has the fastest population rhythm produces more output spikes (***Figure 5***B, green curve). In other words, as WM item strength is varied, it remains the case that whichever PC population in the output gate has the shortest period between volleys of spikes will dominate the output layer, even if each volley contains fewer spikes than the opposing population would otherwise.

**Figure 5.**
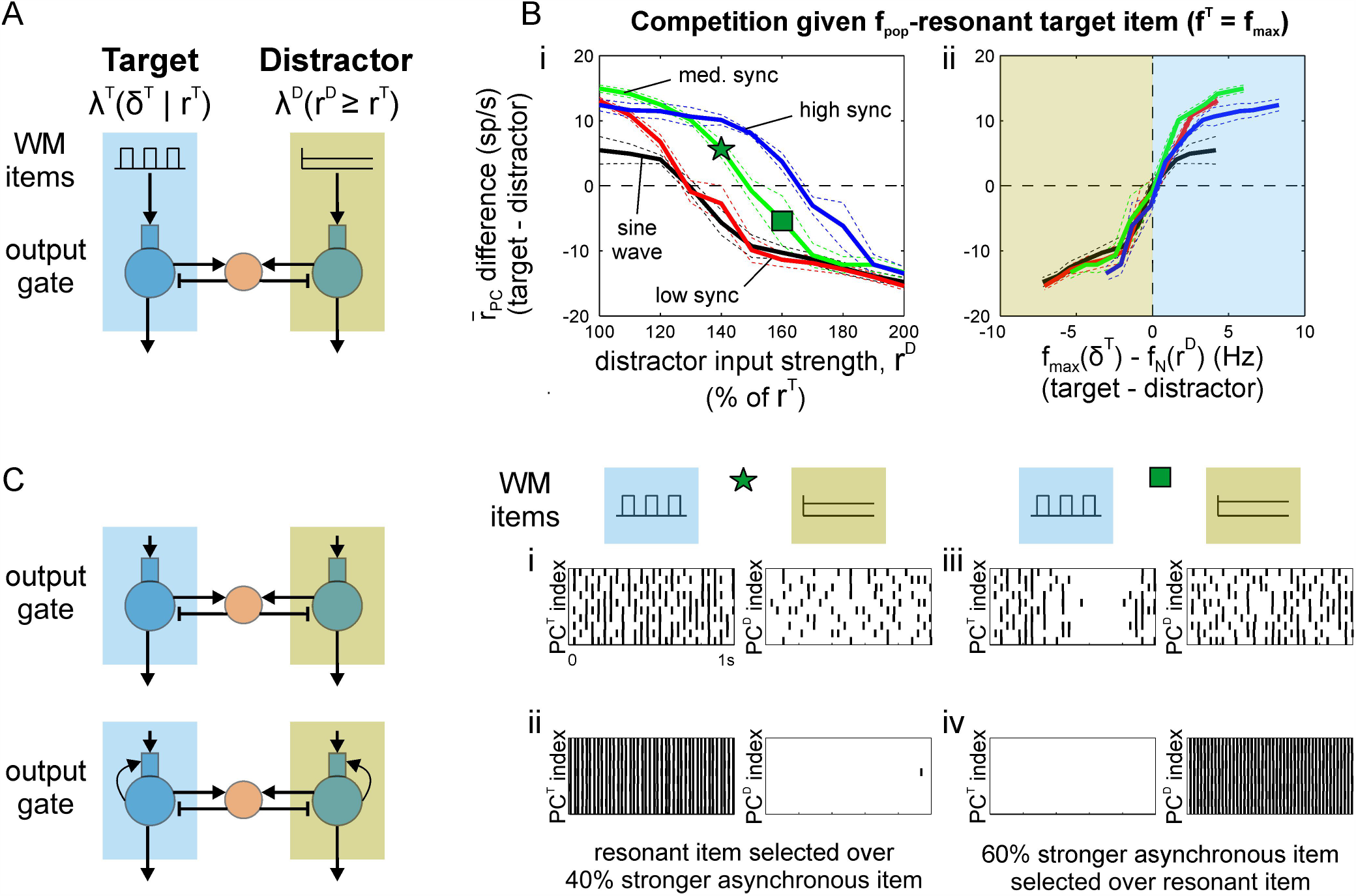
Effect of item synchrony on strength of resonant bias. (**A**) Schematic showing target output (PC^T^) receiving an oscillatory input with synchrony varied across simulations in competition with a distractor output (PC^D^) receiving asynchronous input with strength varied 1-2x the strength of oscillatory input across simulations. Oscillatory inputs were either sinusoidal or square wave with high synchrony (1 ms pulse width), medium synchrony (10 ms pulse width), or low synchrony (19 ms pulse width). (**B**) Differential output firing rates (target-distractor) for target input frequencies maximizing output population frequency (i.e., for inputs at the *f*_*pop*_-resonant frequency) and distractor inputs with increasing strength (i.e., asynchronous input rate, *r^D^*). Differential output is plotted against (i) the strength of distractor input and (ii) the difference in population frequencies expected in the absence of competition (i.e., in an isolated PC/IN network). The blue shaded region highlights the range of responses where target output frequency exceeds the natural frequency expected for the distractor output; the green shaded region highlights the range where distractor output oscillates faster. The star and square in Bi mark distractor strengths that produce greater target and distractor outputs, respectively, and are used to investigate the effects of recurrent excitation. In all cases, the output with higher spike rate was the output with higher population frequency (i.e., a shorter period between spike volleys). (**C**) Recurrent excitation amplifies output differences for winner-take-all selection. (i) Without recurrent excitation, target output is greater despite the distractor receiving a 40% stronger input. This simulation corresponds to the point marked with a star in Bi. (ii) Recurrent excitation amplifies resonant bias producing winner-take-all dynamics that select the output driven by a weaker resonant input. (iii) Without recurrent excitation, distractor output is greater when it receives an asynchronous input that is 60% stronger than an opposing resonant input. This simulation corresponds to the point marked with a square in Bi. (iv) The response to an oscillatory target is suppressed when the stronger asynchronous input elicits a faster natural frequency.

Furthermore, we have shown previously that the *f*_*pop*_-resonant frequency of the output layer depends on the synchrony of its input (***Sherfey et al., 2018a***). That implies that the maximum frequency of the output gate, for a given input strength, can be increased or decreased by varying spike synchrony of an oscillatory item in WM. In other words, the shortest period between volleys of spikes that can be achieved in a rhythmically-driven output population can be decreased by increasing the synchrony of spiking between cells of an input population. This is the case because the ability of inhibitory interneurons in the output gate to spike on every cycle of the excitatory input increases with the synchrony of the feedforward spikes driving them (see ***Sherfey et al.*** (***2018a***) for more details). The consequence of this property for output gating is significant and important because synchrony, itself, is task-modulated during cognitive control tasks (***Buschman et al., 2012***).

Resonant WM items with different degrees of synchrony can induce maximal frequencies in the output gate that exceed the natural frequency to varying degrees. In the high-synchrony case (i.e., all spikes on a given cycle occur within 1 ms), a 31Hz resonant item in WM can produce more spike output than a 70% stronger distractor (***Figure 5***B, blue curve). In the low synchrony (i.e., spikes occurring within 19 ms) and sinusoidal cases, a resonant item can still dominate a 30% stronger distractor (***Figure 5***B, black and red curves). In all cases, the item producing PC spike volleys with the shortest period (i.e., the fastest population rhythm) dominates the output gate (***Figure 5***B, all curves).

### Recurrent excitation amplifies bias for winner-take-all output gating

We have seen that changing target *f*_*pop*_ (by varying *f*^*T*^ and synchrony of the periodic item) or distractor *f*_*pop*_ = *f*_*N*_ (by varying *r^D^* in the asynchronous item) creates a bias in the output gate that favors one item or the other based on the relative population frequencies induced in competing PC outputs (***Figure 2***C, ***Figure 5***B). In the case of large differences between the output population frequencies (e.g., induced by a resonant or a strong asynchronous item), we have seen that this bias can produce a gated response (***Figure 3***, ***Figure 4***). Next, we investigated whether we could achieve winner-take-all (WTA) dynamics by incorporating a common motif in WTA networks.

Specifically, WTA dynamics are commonly observed in networks with a suitable combination of strong lateral inhibition (i.e., inhibition of neighboring excitatory neurons) and strong recurrent excitation (i.e., excitation of neighboring excitatory neurons) (***Kaski and Kohonen, 1994***). Our previous model of the output gate (***Figure 1***B) already contains strong inhibition. When we added recurrent excitation among neighboring cells within each output population, the bias observed previously was, indeed, converted into a WTA response (***Figure 5***C). In the control network, the 28Hz resonant item produced more spiking in the output gate PCs driven by that item than the competing PCs driven by a 40% stronger asynchronous item. With recurrent excitation, the resonant item was selected in the output gate while the asynchronous item was completely blocked. Conversely, when the asynchronous item was made 60% stronger, so that its natural frequency exceeded 28Hz, it was selected in the output gate while the resonant item was completely blocked. The switch from a differential firing rate response to WTA selection across different levels of distractor strength is shown systematically in Supplementary Figure 2. This suggests that strengthening recurrent excitation, potentially through learning input/output mappings (e.g., across trials of a task), can convert a rhythm-mediated bias into an exclusive, WTA output gate that selects the WM item that induces the fastest output oscillation (i.e., the output with the shortest period between spike volleys).

### Multiple rhythmic items in competition

All gating simulations so far have examined the competitive interaction of responses driven by two items, one in an oscillatory state and one in an asynchronous state. However, experiments have shown that multiple rhythms often exist in the same region of cortex (***Lundqvist et al., 2016***; ***Adams et al., 2017***). During WM tasks, both gamma and beta-rhythmic activities have been observed in the same superficial layers of LPFC (***Bastos et al., 2018***). We next asked whether the relative output population frequencies govern competitive dynamics in the presence of multiple rhythmic items in WM. First, we established a reference response based on two rhythms driving separate networks. We then answered the question by examining whether competition induced a deviation that depended on the population frequencies.

The reference case was based on a PC/IN network with a single PC population (***Figure 2***Bi) where PC spiking peaked for *f*^*T*^ ≈ 24 Hz (***Figure 2***Bii, blue curve) while IN spiking and *f*_*pop*_ peaked for *f*^*T*^ ≈ 28 Hz (***Figure 2***Bii-iii) (for medium-synchrony inputs). Given two separate PC/IN networks (i.e., without competition) (***Figure 6***Ai) the relative spike output is determined only by the 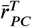 tuning curve in ***Figure 2***Bii. To characterize the reference response, we varied item frequencies over a grid from 16-36 Hz, arbitrarily labeled one the “target” and the other the “distractor”, and then plotted whether target output exceeded distractor output over the (*f*^*T*^, *f*^*D*^) grid (***Figure 6***Aii). This plot shows that the dominant population switches as the input frequencies pass the peak FR at 24 Hz (***Figure 6***Aii, circle). The black zone to the lower-left of the circle in ***Figure 6***Aii indicates that while *f*^*D*^ < *f*^*T*^ < 24 Hz, the target produces more output; however, as *f*^*T*^ increases beyond 24 Hz and *f*^*D*^ remains closer to the 24 Hz input that maximizes spiking, the distractor produces more output. Similarly, the black region to the lower-right indicates that the (distractor) population driven by a higher frequency, *f*^*D*^, produces fewer spikes when the target population is driven by a frequency, *f*^*T*^, closer to 24 Hz. Essentially, as would be expected, whichever PC population in the output gate (without competition) receives an input closer to the frequency (24 Hz) that maximizes PC spiking in ***Figure 2***Bii will output more spikes here. The blue lines mark the higher frequency (28 Hz) at which *f*_*pop*_ peaks in the output (***Figure 2***Biii). If relative population frequencies determined which PC population produced more spike output, we would expect the switch to occur at 28 Hz (i.e., the maximal output frequency) and the target to dominate the region marked with an “x” as well (i.e., 24 Hz < *f*^*D*^ < *f*^*T*^ < 28 Hz).

**Figure 6.**
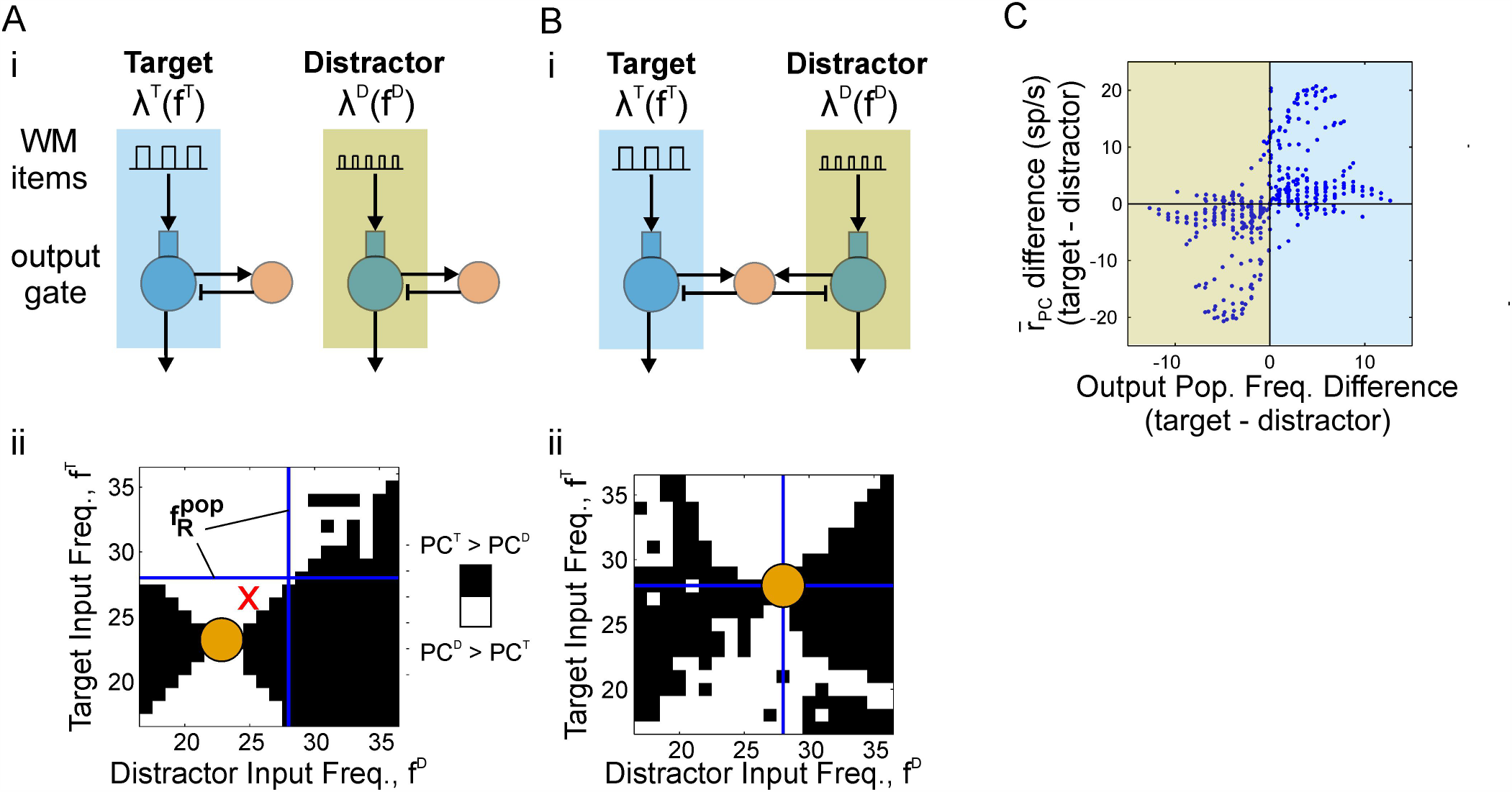
Competition between oscillatory items: the most *f*_*pop*_-resonant item wins. (**A**) Without lateral inhibition, relative spike outputs are determined by 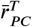 tuning curves in ***Figure 2***Bii. (i) Circuit diagram showing two independent PC/IN networks without lateral inhibition. Both PC populations received oscillatory WM inputs with different modulation frequencies. The modulation frequency of items delivered to each PC population was varied from 18-36 Hz across simulations. Each pathway was arbitrarily labeled “Target” or “Distractor”. (ii) A binary image indicating whether the Target or Distractor PC population outputs more spikes across the simulation given oscillatory inputs at different frequencies. A black pixel indicates the Target population produced more spikes, whereas a white pixel indicates the Distractor population output more spikes. The circle marks the intersection of inputs at the 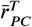 resonant frequency. The blue lines mark the *f*_*pop*_-resonant frequency. Whichever output is closer to peak 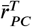 produces more spikes when the circuits are disconnected. (**B**) With lateral inhibition, relative spike outputs are determined by *f*_*pop*_ tuning curves in ***Figure 2***Biii. (i) Circuit diagram showing two PC populations competing through lateral inhibition. The inputs were the same as in (A). (ii) Same plot as (Aii). Whichever output is closer to peak *f*_*pop*_ produces more spikes when the output populations are connected through shared inhibitory interneurons. (**C**) Relative spike outputs with competition plotted against the relative population frequencies without competition. In most cases, the population with higher population frequency produces more spikes.

Indeed, when we have the two outputs compete in the output gate (***Figure 6***Bi), the target most often produces more output as long as *f*^*D*^ < *f*^*T*^ < 28 Hz (***Figure 6***Bii). In this range, the population labeled “target” has the highest population frequency. In contrast to the scenario without competition, whichever population in the output gate (with competition) receives an oscillatory input closer to the maximum population frequency (28 Hz) in ***Figure 2***Biii will produce more spikes here. Furthermore, plotted a different way, we can see that the output with the shortest period between spike volleys usually produces more spikes (***Figure 6***C, lower-left and upper-right quadrants). In contrast to the analogous case for rhythmic vs. asynchronous items (***Figure 5***Bii), however, there are occasions when the lower-frequency population produces more output spikes (***Figure 6***C, upper-left and lower-right quadrants). This occurs when the slower output frequency is closer to the firing rate resonant frequency and the oscillatory input regularly arrives late enough during the inhibitory phase to elicit spikes; however, that is not the case when inputs are weak or low synchrony (not shown).

### Tuning the output gate

A useful gate is one that can be dynamically tuned to select different types of inputs. We have previously shown that the resonant frequency of a PC/IN network can be adjusted by changing the excitability of the PC population via neuromodulation of intrinsic ion channels (***Sherfey et al., 2018a***). Here, we tested whether a similar adjustment of the network’s excitability could be used to flexibly tune an output gate with biased competition for selecting different frequencies of rate-modulation.

In the control case, the network exhibited a maximum output frequency of 28 Hz in response to an oscillatory drive. When driven by items modulated at higher gamma frequencies (e.g., 40 Hz), its output converged to the natural frequency, *f*_*N*_ ≈ 21 Hz (***Figure 2***Biii). In contrast, the model output frequency equaled the input frequency below 28 Hz. Thus, in the present model (***Figure 1***B), a 25 Hz (i.e., < *f*_*max*_) input rhythm induces a higher output frequency than a higher 40 Hz (i.e., < *f*_*max*_) input rhythm. Consequently, biased competition in the output gate favors the beta-rhythmic item and blocks responses to the gamma-rhythmic item (***Figure 7***A). How can we selectively output the gamma-rhythmic item?

**Figure 7.**
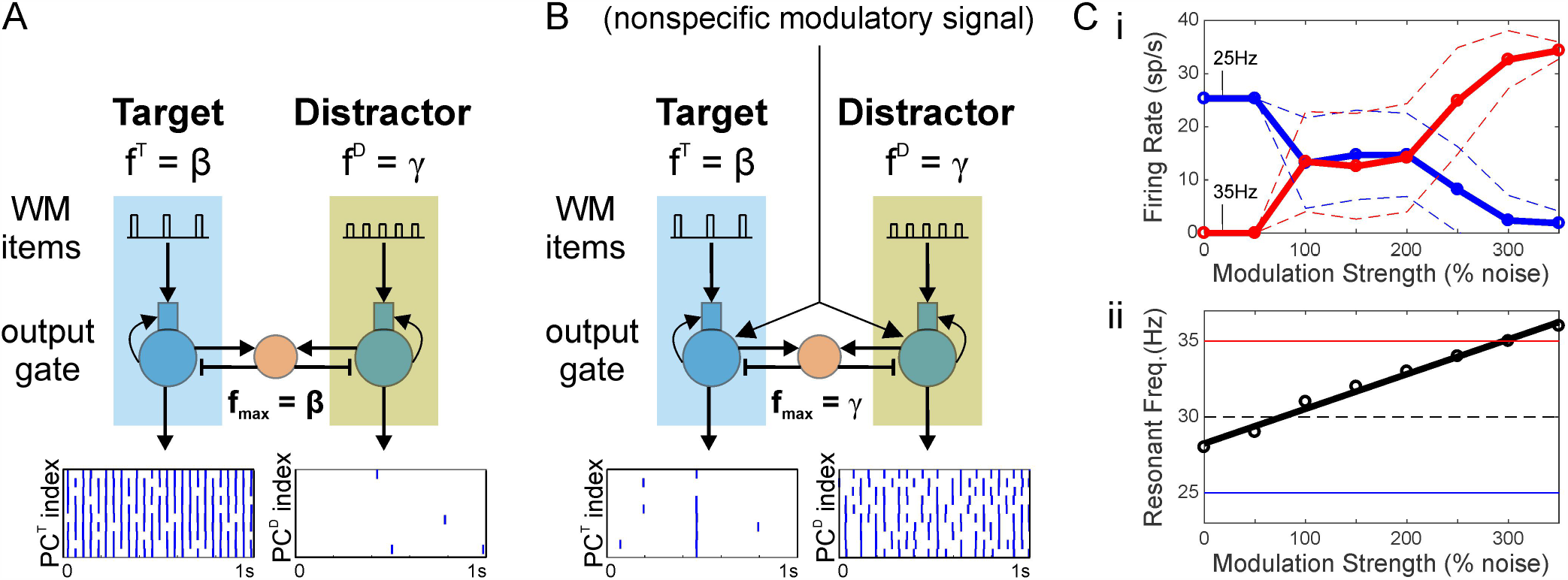
Nonspecific inputs can tune output resonance for switching between specific beta- and gamma-rhythmic pathways. (**A**) A resonant beta input suppresses the response to a less resonant gamma-frequency input. (**B**) A nonspecific asynchronous input to both output populations shifts their resonant frequency to the gamma-range, causing the output layer to select the gamma-rhythmic input and suppress response to the less resonant beta input. (**C**) Tuning the network by varying modulation strength. (i) Firing rate of two, competing PC populations: one driven by a 25Hz oscillation (blue) and the other driven by a 35Hz oscillation (red). Solid and dashed lines show the mean and mean ± standard deviation, respectively. The strength of a nonspecific, modulatory (asynchronous) signal to both populations is increased from 0% to 350% of the background spiking. Without the signal, the 25Hz signal is exclusively propagated (i.e., always wins the competition). With strong modulation, the 35Hz signal is exclusively propagated. For intermediate levels of modulation, there may be spiking in either population at different points in time. (ii) Resonant frequency of the output gate plotted against nonspecific modulation to PCs of the gate. The thick solid line shows a linear fit. The thin horizontal lines mark the two input frequencies (solid) and the midpoint between them (dashed).

Building on our earlier work (***Sherfey et al., 2018a***), we hypothesized that increasing the excitability of the output gate would increase the gate’s resonant frequency and lead to the selection of WM items with a higher modulation frequency. To test this, we delivered a nonspecific, subthreshold, asynchronous drive to all PC populations of the output gate, effectively increasing their resting membrane potential and, thus, their excitability. With sufficient strength, such a signal enabled the output gate to follow the gamma-rhythmic item. The response to the gamma-rhythmic item then exhibited a shorter period between volleys of inhibition-recruiting spikes and, consequently, suppressed the response to the beta-rhythmic item (***Figure 7***B). Thus, the gamma-rhythmic item was exclusively selected by making the output gate resonate to gamma frequencies. As modulation strength increases, the resonant frequency increases smoothly and switches between the exclusive propagation of lower frequency signals to propagation of higher frequency signals (***Figure 7***C). This represents a flexible mechanism by which a nonspecific modulatory signal can be adjusted to tune the maximum frequency of the output gate so that it selectively responds to items modulated at variable target frequencies.

### Oscillatory gating for population-coded items

The WM items we have investigated systematically throughout this work had time-averaged firing rates that were uniform across the input population. However, experiments have shown that PFC encodes items in the pattern of firing rates across PCs of an encoding population (***Mante et al., 2013***). For our oscillatory gating mechanism to be used to gate outputs from WM, it must be able to gate such population rate-coded items. Next, we tested the ability of our oscillatory gating mechanism to select rate-coded items with partial Gaussian spatial profiles (***Figure 8***). These examples demonstrate the utility of our gating mechanism for propagating more complex signals and its ability to operate in feedforward pathways with both parallel and convergent projections.

**Figure 8.**
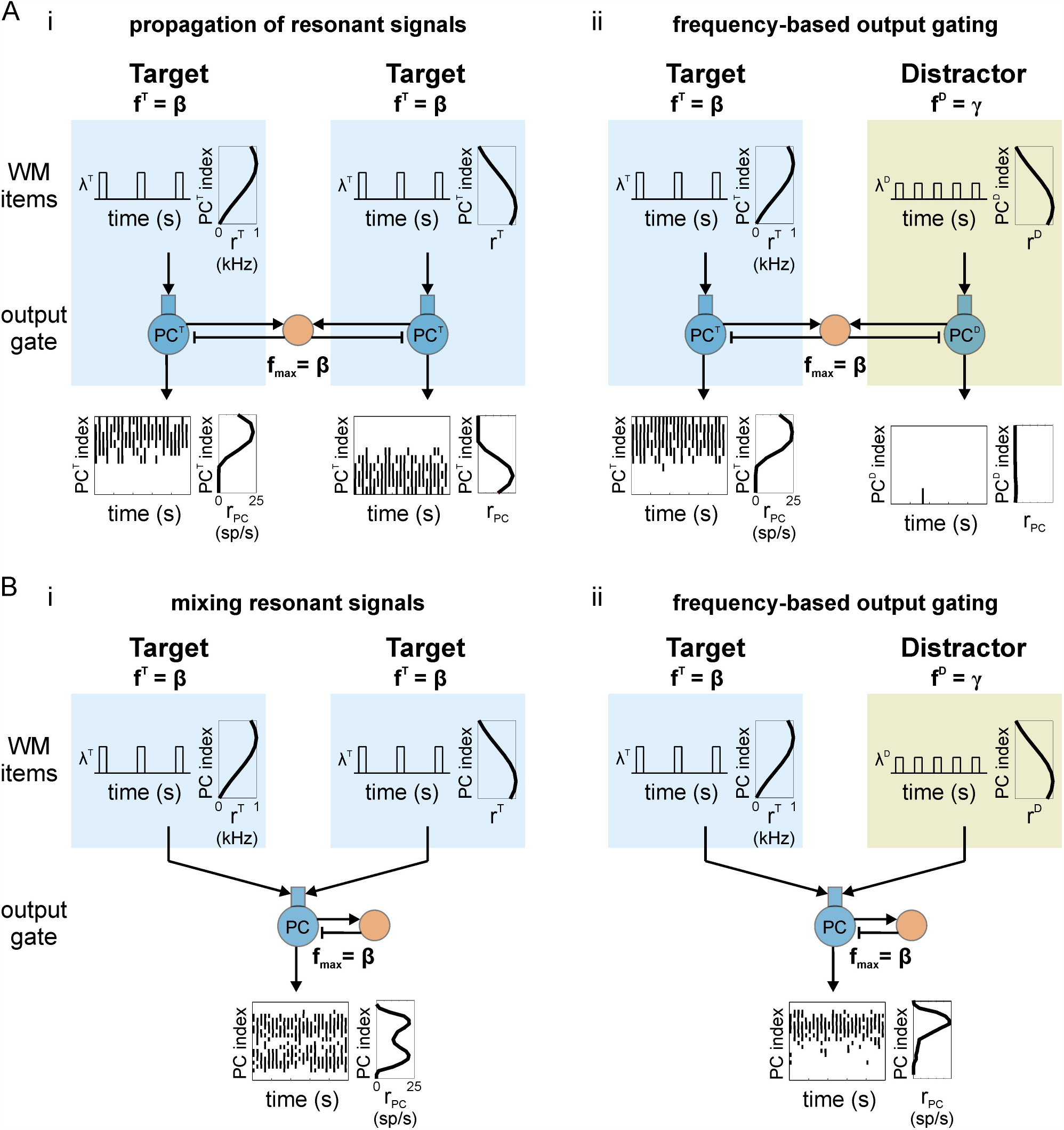
Oscillatory gating for population-coded signals. (**A**) Resonant bias supports frequency-based gating of population rate-coded signals among parallel pathways. (i) Outputs of two pathways reflect the spatial pattern of firing rates across their inputs when both inputs are embedded in resonant oscillations. PC index represents linear indices of cells in a given output PC population. The spatial pattern of time-averaged firing rates across cells of the population is assumed to encode item information. Individual cells have low firing rates while the collective population is modulated at a faster frequency. Both WM items have the same modulation frequency. (ii) Frequency-based output gating: More resonant rate-coded signals suppress less-resonant rate-coded signals. WM items have the same spatial pattern of time-averaged firing rates as in (Ai), but now the frequency of faster time scale modulation differs between the two items; only the Target item has a modulation frequency that is resonant with the output gate. (**B**) Resonant bias supports frequency-based gating of rate-coded signals among convergent pathways. (i) Similar to (A) except both inputs converge on a single output population that reflects the approximate sum of the input signals in its spatial pattern of firing rates. (ii) A less resonant gamma-frequency signal is blocked from the output population.

#### Parallel pathways

FIG8First, we simulated two items with resonant beta-frequency modulation of the instantaneous firing rate and different Gaussian spatial patterns of time-averaged firing rates. When delivered to two PC populations in the output gate, the spatial pattern was largely conserved in the corresponding PC populations (***Figure 8***Ai). This demonstrates that parallel, rate-coded items that are resonant with the output gate can be propagated simultaneously. In contrast, when the fast modulation of the instantaneous firing rate occurs at a non-resonant gamma-frequency, only the resonant beta-frequency item is conserved in its output population while the non-resonant item is blocked (***Figure 8***Aii). This represents a form of frequency-based output gating that selects for the most resonant rate-coded item in WM.

#### Convergent pathways

Anatomically, projections from superficial to deep layers of cortex involve some degree of overlap (***Levitt et al., 1993***; ***Kritzer and Goldman-Rakic, 1995***) and this, most likely, results in multiple WM items projecting to some of the same cells in the deep layer. We have shown throughout this paper that lateral inhibition driven by a resonant output gives rise to periodic inhibition that blocks responses in opposing pathways. Next we tested the hypothesis that feedback inhibition can similarly block responses to less resonant rate-coded signals when pathways converge on a single output population. Similar filtering has been demonstrated in simpler models with uniform input rates (***Cannon et al., 2014***).

We repeated the previous simulations demonstrating frequency-based gating of rate-coded signals, except this time we delivered both items to the same PC population in the output gate (***Figure 8***B). In this case, two rate-coded items with Gaussian profiles and resonant beta-frequency modulation were approximately conserved in the spatial pattern of output firing rates in the single PC population (***Figure 8***Bi). In contrast, when one item was modulated at a less resonant gamma frequency, the item with resonant modulation was conserved in the output while the response to the less resonant item was blocked by the feedback inhibition as hypothesized (***Figure 8***Bii).

## Discussion

We have presented a novel mechanism for gating the propagation of signals through a relay network and illustrated its utility for gating outputs from a working memory (WM) buffer. The mechanism requires strong feedback inhibition in the output gate to induce periodic responses. Essentially, the output population with the shortest period between spike volleys will most reliably engage inhibitory interneurons to suppress responses in competing outputs. Resonant properties that result from the feedback inhibition support frequency-based output selection of items in the WM buffer. This enables the modulation frequency of WM items to govern output selection, possibly in opposition to the strengths (i.e., time-averaged firing rates) of items in WM. We showed that the resonant frequency can be tuned by flexible modulatory signals to select different frequencies. We also showed that with the addition of recurrent excitation (e.g., with learning), the gate can exhibit exclusive, winner-take-all responses. Finally, the same oscillatory gating mechanism could be used to gate flows, in general, for arbitrary, population rate-coded signals along parallel and convergent pathways in any cortical region.

### Summary of oscillatory gating rules

The results from this study can be organized into a set of rules that depend on dynamical states and determine which WM item(s) will dominate the output gate.

#### Asynchronous vs. asynchronous signal

The strongest asynchronous signal will produce the strongest response in the output gate because it induces the highest natural frequency (i.e., the shortest period between spike volleys) (see ***Figure 2***A for the natural response to asynchronous items and ***Figure 4***, lower plots, for dependence of natural frequency on item strength).

#### Beta/gamma rhythmic signal vs. asynchronous signals

A (possibly weaker) rhythmic input will produce the strongest response in the output gate if it induces a population response with the shortest period between spike volleys (***Figure 3***, ***Figure 4***A). That occurs when the modulation frequency of the input signal is between the strength-dependent natural frequency induced by the asynchronous signals and the input synchrony-dependent maximum frequency of the output (***Figure 5***B, Ci-ii). Otherwise, the asynchronous signal can produce an equally-strong or stronger response (***Figure 4***B, ***Figure 5***Ciii-iv).

#### Rhythmic vs. rhythmic signals

The input with modulation frequency that is most resonant (***Figure 6***) with the tunable output gate (***Figure 7***) will produce the strongest response.

### Control mechanisms

According to the gating rules, responses in the output gate depend on the resonant frequency of the gate (i.e., its maximum frequency) as well as the synchrony and modulation frequency of rhythmic items in the input layer. Importantly, tuning relay network resonance can flexibly guide the flow of information from input populations without requiring changes in the relative strengths of population activity. The resonant frequency can be rapidly tuned by changing the level of background excitation present in the output gate using an asynchronous modulatory signal (***Figure 7***), possibly originating in thalamus or the deep layers of another region of PFC (***Barbas, 2013***).

### Oscillatory gating for cognitive function

#### Working memory

In the brain, WM tasks have implicated the prefrontal cortex (PFC) in all aspects of WM and cognitive control (***Fuster, 1973***, ***2015***; ***Goldman-Rakic, 1995***; ***Miller, 2000***). We hypothesize that the deep layers of PFC function as an output gate for WM items represented in a superficial buffer. Activation of select populations in the output gate would determine when WM items influence downstream circuits and which downstream circuits (i.e., systems and processes) are influenced (***Ardid et al., 2007***; ***Kriete and Noelle, 2011***; ***Badre and Frank, 2012***; ***Kriete et al., 2013***; ***Ardid and Wang, 2013***; ***Hasselmo and Stern, 2018***; ***Zhu et al., 2018***, ***2020***).

Task-relevant WM items, presumably engaged in ongoing processing, often exhibit synchrony at beta and gamma frequencies (***Siegel et al., 2009***; ***Lundqvist et al., 2016***, ***2018***; ***Bastos et al., 2018***). Our findings suggest that such resonant oscillations could support the selective output of WM items even when other less-resonant items are stored with stronger activation (e.g., items stored in asynchronous population-coded signals with 70% higher mean firing rates) (***Figure 5***). The selected items would then be available in an oscillatory state for read-out in subcortical structures and participation in downstream processing. Consistent with this hypothesized mechanism for dynamic routing of WM representations, beta-synchrony has been observed between PFC and higher-order thalamus during a WM task, and it was correlated with performance (***Parnaudeau et al., 2013***).

Our model predicts that oscillation-based dynamic routing would produce elevated phase locking between the WM item-encoding neurons in the superficial buffer and the deep layer output gate during WM retrieval. The model also predicts that the resonant frequency of the deep layer network will be tuned to the population frequency exhibited by target item-encoding superficial populations. This implies that altering the excitation of the deep layers, even globally, will impact the coherence between superficial and deep layers, specifically in regions associated with a particular target input. If a correct response requires a certain resonant frequency in the deep layers, the model implies that spontaneous mistakes may show up as changes in the excitability (and hence resonant frequency) of the deep layers. Furthermore, an experimental change in excitability of the deep layers may increase error rates. These are testable hypotheses.

#### Rule-based action and cognitive control

Context-dependent changes in the oscillatory state of WM items or the resonant frequency of the output gate could mediate rule-based selection of input-output mappings. Experiments have revealed that populations coding task-specific rules exhibit increased beta-rhythmic synchrony (***Buschman et al., 2012***). In our model, the deep layer output gate exhibits resonance at a similar frequency and consequently supports the selective read-out of WM items that are synchronous with beta-frequency oscillations. Rule-dependent synchrony of WM items, then, could control which input-output mappings are engaged while suppressing responses of opposing outputs via lateral inhibition in the output gate (see ***Figure 3***). Rule updating via oscillatory state control mechanisms operating on the WM buffer could potentially be directed by contextual inputs from hippocampus (***Komorowski et al., 2013***), error signals from ACC (***Kerns et al., 2004***; ***Amiez et al., 2005***), or basal ganglia gating mechanisms (***Frank et al., 2001***; ***Badre and Frank, 2012***). Our model predicts that experimental perturbations that facilitate or disrupt the control of oscillatory states could improve or degrade context specificity of response.

Furthermore, a PFC output gate could govern flows along parallel pathways from visual and auditory cortices (***Barbas, 2015***) through subregions of basal ganglia and thalamus (***O’Reilly and Frank, 2006***), effectively sculpting functional connectivity between disparate regions. For instance, the oscillatory gating mechanism could dynamically link conditions encoded in sensory cortices to subcortically-mediated responses that guide attention, action, and cognitive control, more broadly. Thus, the mechanism may contribute to dynamics engaged by many cognitive processes.

### Comparison to other work

Theoretical work has shown that input gating mechanisms, controlling whether WM items are maintained or updated, can effectively manage a tradeoff between stability and flexibility of the WM (***Hochreiter and Schmidhuber, 1997***; ***Miyake and Shah, 1999***; ***O’Reilly and Frank, 2006***; ***Badre and Frank, 2012***) and that output gating mechanisms provide additional control over when maintained WM items effectively influence downstream targets (***Hochreiter and Schmidhuber, 1997***; ***Kriete and Noelle, 2011***; ***Kriete et al., 2013***). Experiments suggest input and output gating of WM items are present in PFC (***Badre and Frank, 2012***), while models of input (***O’Reilly and Frank, 2006***) and output (***Kriete and Noelle, 2011***; ***Hasselmo and Stern, 2018***; ***Zhu et al., 2018***, ***2020***) gating of WM items in PFC have proposed activity-based mechanisms mediated by interactions between PFC and basal ganglia (BG). We have presented an alternative gating mechanism that is mediated by known network oscillations within cortex, instead of subcortical activity levels. In contrast to activity-based mechanisms, oscillatory gating enables the flow of information to be governed by more flexible and rapid alterations in spike timing instead of more slowly-changing activity levels (i.e., spike rates).

***Frank and Badre*** (***2012***) have shown how the basal ganglia can gate outputs from a PFC WM buffer using an activity-based, disinhibitory mechanism. In their model, BG regulates a signal that increases or decreases thalamic activity to regulate whether PFC outputs can be sufficiently activated for read-out. It is possible that their activity-based mechanism operates in parallel with our oscillation-based mechanism or that the two work in conjunction. For instance, BG-regulated thalamocortical activity could provide an asynchronous modulatory signal that tunes the resonant frequency of an oscillatory output gate in PFC. Given rule-specific BG activity, this conjunctive approach could represent a novel mechanism for selecting rule-based input-output mappings using a combination of activity- and oscillation-based gating.

### Conclusions

In this work, we introduced a new oscillatory gating mechanism that enables beta- and gamma-frequency oscillations to govern read-out from working memory. The output gate can by rapidly tuned to support routing for flexible cognitive processes; the same mechanism leveraging fast network oscillations and local inhibition can shape functional connectivity throughout neocortex.

## Supplementary Material

**Supplementary Figure 1.**
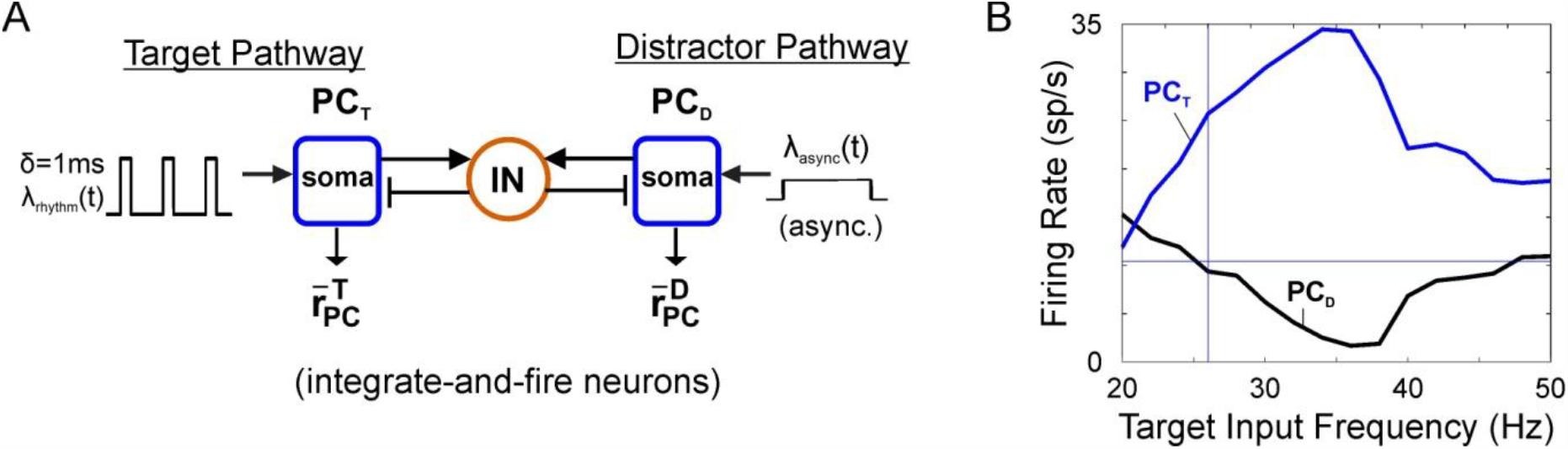
Competitive interaction between populations in a leaky integrate-and-fire (LIF) network. (A) Schematic showing asynchronous Poisson spike trains driving principal cells (PCs) coupled to interneurons (INs) providing strong feedback inhibition. Inputs to the LIF network were the same as the more detailed PFC network described in the Methods section except that g_inp_ = 0.00375 mS/cm^2^ and g_noise_ = 0.0056 mS/cm^2^. See (Sherfey et al., 2018a, S3 Fig) for LIF model equations. (B) Mean firing rate outputs for target (blue) and distractor (red) as target input frequency is increased (compare to Figure 3Cii).

**Supplementary Figure 2.**
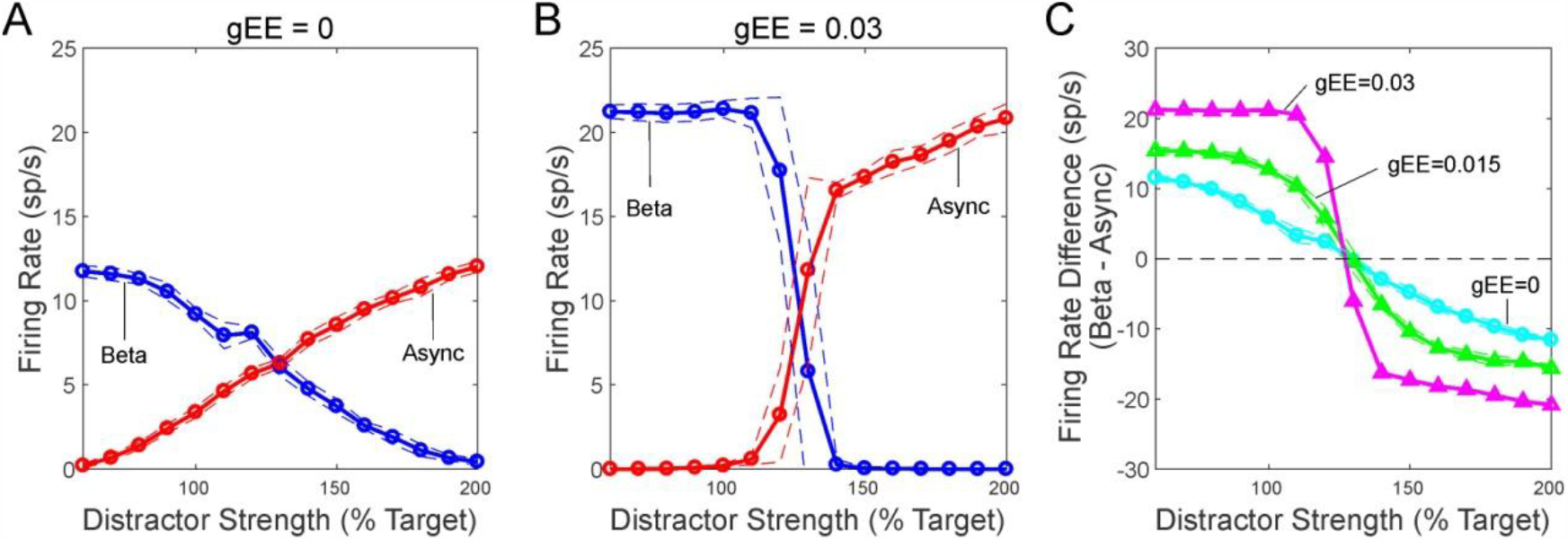
Winner-take-all behavior of the PC/IN network with recurrent excitation. (A) Simulation without recurrent excitation. The plot shows the firing rate of two, competing PC populations: one driven by a resonant, beta2-frequency oscillation (blue) and the other driven by an asynchronous signal with strength that varied from 100% to 200% that of the oscillatory signal. (B) Simulation with recurrent excitation. (C) Comparison of the differential firing rates (similar to Figure 6Bi) between the populations driven by oscillatory versus asynchronous signals. Intermediate levels of recurrent excitation (green) produce a transition between the biased response (cyan) and winner-take-all dynamics (magenta).

